# Quantifying The Impact of Bulk TCR-Seq Methodological Choices on The Profiled T Cell Repertoire

**DOI:** 10.1101/2025.08.05.668736

**Authors:** Aya K.H. Mahdy, Janina Fuß, Sören Franzenburg, Melanie Vollstedt, Yewgenia Dolshanskaya, Érika Endo Kokubun, Valeriia Kriukova, Florian Tran, Corinna Bang, Marte Lie Høivik, Jurgita Skieceviciene, Juozas Kupčinskas, Mathilde Poyet, Andre Franke, Hesham ElAbd

**Author notes:** Shared senior authors (Andre Franke, Hesham ElAbd).

## Abstract

Bulk T cell repertoire profiling using sequencing (TCR-Seq) is a powerful method to investigate T cell responses to natural infections, vaccines, cancers, and autoimmune diseases. This assay can be conducted using various techniques, such as multiplex PCR or 5’-RACE. However, each methods introduces systematic biases that can result in different pictures of the underlying T cell repertoire. Furthermore, the impact of technical variables on the accuracy of these methods remains understudied. Thus, in this study, we systematically characterized different multiplex PCR-based protocols, focusing on the quality and quantity of the utilized RNA/DNA, extraction methods, amplification programs, variations between production batches, and technical handling of samples. Our findings highlight the important role of RNA/DNA quality in shaping the profiling results of T cell repertoires. Whereas low RNA/DNA quantities can be partially compensated for by increasing the number of PCR cycles, this is partially not possible with lower quality. In conclusion, our results highlight the influence of different technical choices on the biological conclusions drawn from TCR-Seq data and provide practical guidelines to finetune these variables to ensure consistent and reliable results under diverse experimental constraints.

## Introduction

T cells are a versatile group of cells that are heavily involved in multiple biological processes ranging from responding to infections^1^, to fighting cancer cells^2^ and mediating tissue hemostasis^3,4^. Based on the sequence of T cell receptors (TCRs), T cells are arranged into clones with the members of a clone sharing the same receptor. These receptors are generated via V(D)J recombination, which generates an extraordinary degree of variation that enables T cells to recognize a wide range of antigens^5^. T cells can be broadly divided into two main categories based on the antigen binding chains used in their T cell receptor, first, alpha/beta (αβ) T cells, which use the TCR-α and TCR-β chains, and second gamma/delta (γδ) T cells, which utilize TCR-γ and TCR-δ chains^6^.

A commonly used method to investigate the collection of T cell clones in biological samples, *i.e.,* the T cell repertoire, is bulk <u>T c</u>ell receptor <u>seq</u>uencing (TCR-Seq). In this method, usually, a collection of primers targeting the different V and J genes that form the individual antigenbinding chains of a TCR, *e.g.,* TCR-α or TCR-γ, is used to amplify the V(D)J recombination products that encode the different TCRs present in a sample. Subsequently, the amplified products are sequenced using <u>n</u>ext-<u>g</u>eneration <u>s</u>equencing (NGS) and processed using various computational workflows to generate a list of recombination products, as well as quantification of their relative expansion-*i.e.* an estimate of their clonal size^7^.

Different technical and biological reasons hinder the accurate identification and quantification of the T cell clonotypes present in a sample, such as the sensitivity of the assay to detect rare clonotypes, amplification biases introduced during the PCR reaction, sequencing errors, multiplexing strategy, *et cetera*. Additionally, the starting biological material is generally a subset of the repertoire, *e.g.* a few milliliters of blood. Although TCR-Seq is a commonly used protocol, fewer studies^8,9^ have investigated the influence of different methodological choices on the profiled T cell repertoire. For example, Barennes *et al.^9^* reported significant systematics biases among different repertoire profiling methods. While these studies such as the Barennes *et al.^9^*, compared the results of different amplification protocols, which is valuable for selecting an appropriate technique for a given experimental design, they do not provide a guideline for navigating the complex set of technical variables that need to be fine-tuned in each assay.

To address this problem, we here investigate the impact of different technical variables on two commercially available TCR-Seq protocols. We started by examining the impact of amplification strategy, specifically in terms of PCR cycles, and the quantity and quality of DNA or RNA used. Thereafter, we explored technical factors influencing the profiling of the T cell repertoire of different tissues, namely, the blood and the colon. Lastly, we investigated the impact of the RNA extraction method, the impact of different production lots, and the technical handling of samples.

## Results

### RNA quantity and quality heavily influence the detection of T cell clonotypes

As a first step, we isolated RNA from PAXgene tubes collected from eight different donors. The isolated RNA varied in quality (**Supp. Table S1**) as measured by the RNA integrity number equivalent (RIN^e^) score, ranging from 4 to 9.3 with a mean and a median of ~7. Subsequently, the αβ T cell repertoire of these samples was measured using a commercially available TCRSeq library preparation kit according to the manufacturer’s instructions (**Material and Methods**). From each sample, three libraries were prepared using different amounts of isolated RNA, namely, 100, 200, and 300 ng, using the same PCR amplification program (18 cycles for PCR1 and 15 cycles for PCR2). After sequencing the generated libraries using NGS (**Material and Methods),** MiXCR^7^ was used to identify the α and β T cell repertoire from each prepared library in compliance with the manufacturer’s recommendations (**Material and Methods)**.

Initially, we started by comparing the number of reads identified from the libraries defined above. Across all samples, libraries prepared with different RNA quantities yielded different numbers of total sequencing reads (**Fig. S1A-H**). Subsequently, we processed the quality control report of MiXCR^7^ and extracted the number of reads that aligned to immune gene loci (aligned reads hereafter), as well as the number of reads that failed to align (unaligned reads hereafter). Across all samples, most of the generated reads aligned to the different immune loci (**Fig. S2A-H**) with two notable exceptions, sample 2 (**Fig. S2B**) and sample 6 (**Fig.S2F**), which had a substantial fraction of unaligned reads, *e.g.* in sample 6, the fraction of unaligned reads was higher than aligned reads. These reads were not aligned by MiXCR because they lack barcodes, which is most likely caused by the high adaptor sequences observed in these samples. After that, we wanted to investigate the correlation between increasing the amount of RNA used for library preparation and the number of aligned and unaligned reads. Increasing the amount of input RNA positively correlated with the number of aligned reads (**Fig. S3A**), but not with the number of unaligned reads (**Fig. S3B**) and as a result the ratio of unalignedto-aligned was negatively correlated with the amount of RNA used for preparing the library (**Fig. S3C**). Additionally, the quality of RNA positively correlated with the number of aligned (**Fig. S3D**) and unaligned reads (**Fig. S3E**) and consequently the ratio of aligned-to-unaligned reads (**Fig. S3F**).

After investigating factors impacting the number of reads per sample, we started analyzed clonotypes identified by MiXCR^7^. Across the 48 libraries (two chains x eight samples x three RNA amounts), we obtained a median of 35,146 and 41,447 clonotypes from the repertoire of the α and β chains, respectively. Unsurprisingly, increasing the amount of RNA used for library preparation correlated positively with increasing the number of identified clonotypes for both α and β chains (**Fig. 1A-1B**). Similarly, the quality of RNA influenced the number of identified clonotypes from the repertoire of both chains (**Fig. 1C-1D**). Importantly, increasing the quantity of RNA could not compensate for the lower quality of the RNA across both measured loci (**Fig. 1E-1F**). Specifically, samples with very low-quality scores (RINe score <5) had a much smaller measured repertoire in comparison to samples with a higher-quality RNA even after increasing the amount of RNA used to construct the assay. For example, the repertoire of samples having high-quality RNA (RINe >8) at 200 ng of input RNA had a higher clonal richness, *i.e.* contained a higher number of clonotypes, than the repertoire of samples having low-quality RNA (RINe score <5) at 300 ng of input RNA (**Fig. 1E-1F**).

**Figure 1:**
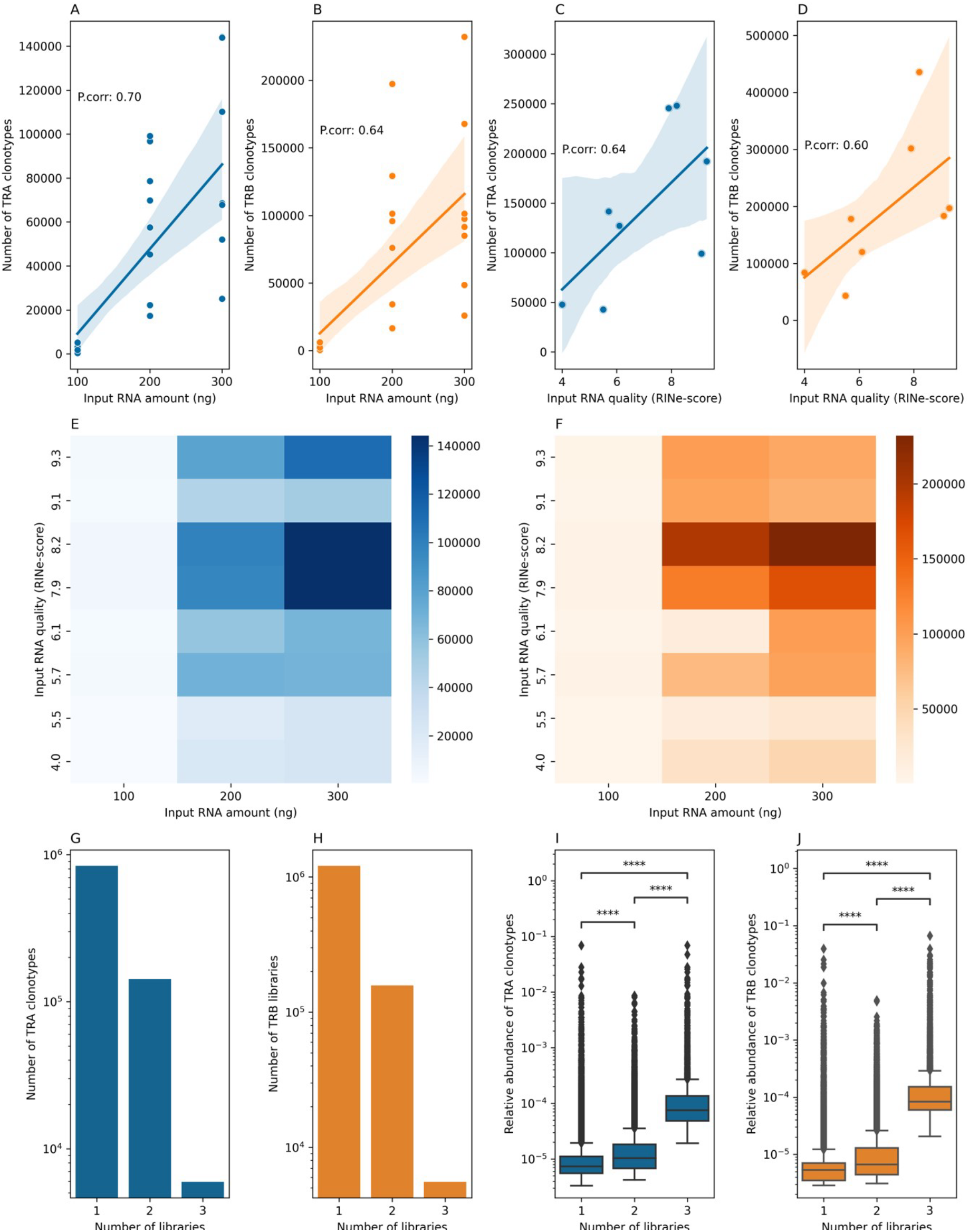
factors governing the ability of TCR-Seq to infer the composition and the structure of the TCR repertoire. **(A)** the relationship between the amount of input RNA used for preparing TCR libraries and the number of identified T cell α clonotypes. The same relationship is depicted in **(B)** but for the number of identified T cell β clonotypes. **(C)** and **(D)** shows the relationship between the RNA quality and the number of identified α and β T cell clonotypes. **(E)** and **(F)** illustrate the impact of RNA quantity and quality on the number of identified clonotypes from the α and β chains, respectively. **(G)** and **(H)** show the number of times a TCR-α **(G)** or TCR-β **(H)** clonotype was identified across the libraries prepared from each sample using three different concentrations, namely, 100, 200, and 300 ng. **(I)** and **(J)** depicts the relationship between the average relative abundance of a TCR-α **(I)** or TCR-β **(J)** clonotype and the number of times it was detected across the three libraries prepared from different RNA concentrations.

After that, we looked at the overlap in clonotypes between the three libraries prepared from each sample using three different RNA quantities to identify factors governing the reproducibility and detectability of the identified clonotypes. For clonotypes derived from the TCR-α and the TCR-β chain, the majority of clonotypes were observed only once and only a small minority of clonotypes were observed among the three replicates (**Fig. 1G-1H**). This minority of TCR-α and TCR-β clonotypes were significantly more expanded than other groups (**Fig. 1I-1J**). This suggests that highly expanded clonotypes can be detected even with a small amount of RNA material, *e.g.* 100 ng in our case, or with low RNA quality, however, the detection of rarer or less expanded clonotypes will be much harder under these conditions.

To follow this further, we examined the behavior of the top 10 most expanded clonotypes at the low amount of input RNA level, *i.e.* 100 ng when increasing the starting material to 200 ng and 300 ng (**Fig. S4**). The cumulative sum of this set of top 10 most expanded clonotypes contributed a substantial fraction of the measured repertoire, in terms of the number and the fraction of unique molecular identifiers (UMI) they occupy (**Fig. S4A-S4B**) and also the count and fraction of reads derived from them (**Fig. S4C-S4D**). This behavior was consistent across the three different quantities of RNA used namely, 100, 200, and 300 ng. These results indicate an interplay between the detection limit of the assay and the input amount of RNA, where at low input, *e.g.* 100 ng, the assay only picks the most expanded clonotypes and it fails to detect other clonotypes. However, with increasing the input amount, the abundance of amplicons derived from other clonotypes crosses the detection limit of the assay (at the utilized sequencing depth) and we start observing other clonotypes (**Fig. 1A-1B**). Nonetheless, the expanded clonotypes still represent a substantial fraction of the reads and UMI because of their abundance, *i.e.* increasing the amount of RNA leads to an increase in the number of unique clonotypes measured, however, most of the reads and UMIs are used by most expanded clonotypes (**Fig. S1)**. Thus, by increasing the amount of input RNA, the sequencing depth needs to be increased to identify rare clonotypes as a substantial fraction of reads and UMI will be consumed by highly expanded clonotypes.

### Optimizing the amplification strategy increases the sensitivity of TCR-Seq in samples with limited input

After that, we wanted to investigate the impact of increasing the number of PCR cycles on the measured repertoire using TCR-Seq. To this end, we used the same experimental design defined above and increased PCR 1 to 21 and 24 cycles and the second PCR reaction to 18 and 21 cycles to have three libraries prepared from the same amount of RNA. At the lowest amount of RNA tested in the experiment, *i.e.* 100 ng, changing the PCR amplification program had the highest impact (**Fig. 2A-2B**) whereas at 300 ng the PCR amplification program had little to no impact on the number of inferred clonotypes (**Fig. 2A-2B**). To make sure that the results are not confounded by artifacts introduced during the PCR reaction, we utilized the UMIs introduced during library preparation and filtered clonotypes with less than two reads (**Fig. S5A-S5B**), as well as clonotypes with less than two different UMIs (**Fig. S5C-S5D**). Although these filtration steps changed the number of identified clonotypes, especially removing clonotypes with less than two UMIs (**Fig. S5C-S5D**), the impact of changing PCR cycles was much more prominent at 100 ng than at 300 ng. Furthermore, we did not observe any skew in the distribution of the number of UMIs per clonotypes among the three different amplification programs, specifically, the majority of clonotypes were supported by less than ten UMI (**Fig. S6A-6B**) with one UMI per clonotype being the median and the mode (**Fig. S6CS6D**). These findings illustrate that increasing the number of PCR cycles did not alter or change the distribution of UMIs per clonotype. Thus, for low quantities of RNA, increasing PCR cycles is a promising strategy to increase the number of identified clonotypes from the samples, however, this approach will have a minimal effect on samples with high quantities of RNA.

**Figure 2.**
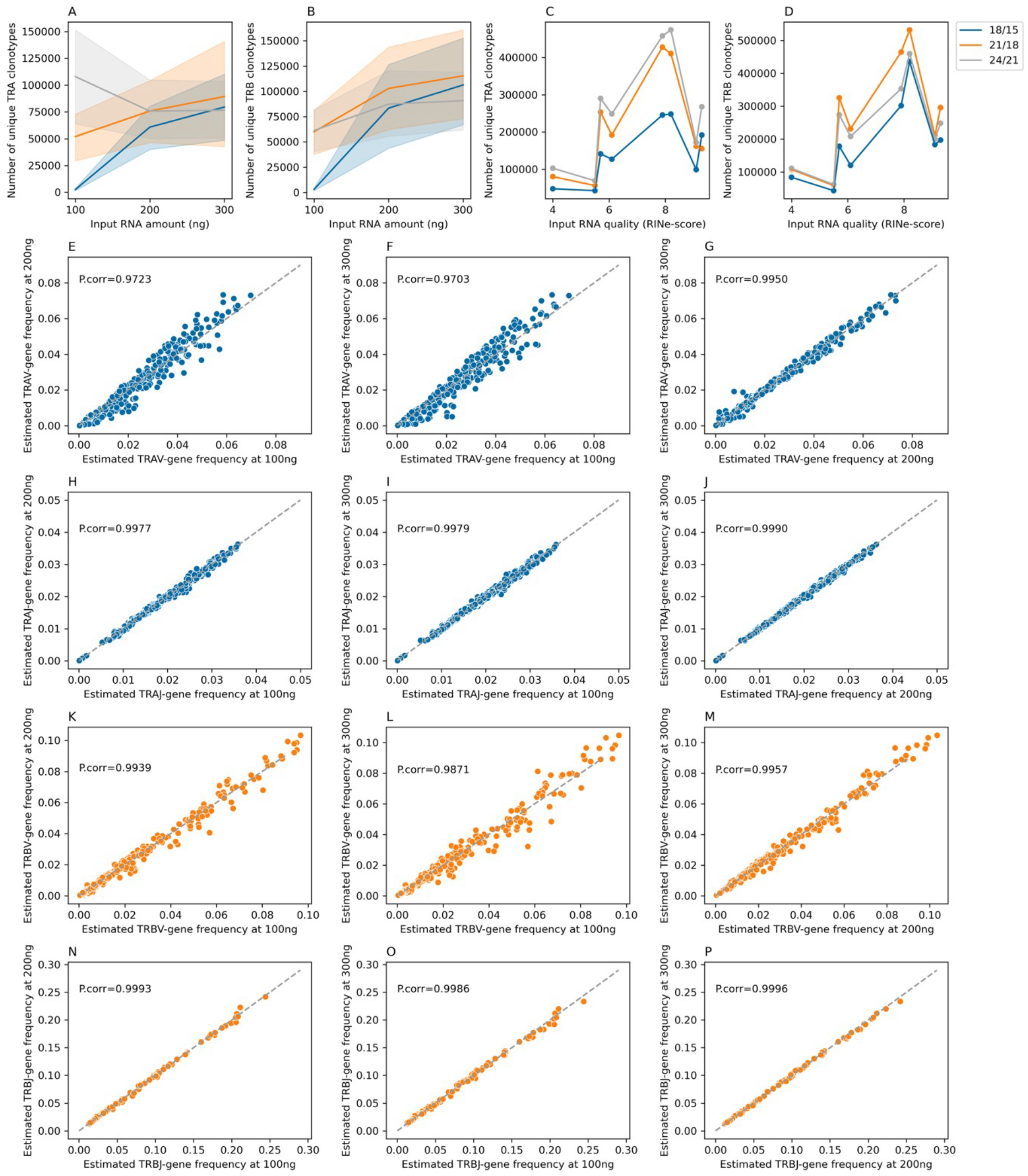
The relation between the amount of input RNA, PCR-cycle optimization, and the number of identified clonotypes. **(A)** and **(B)** illustrate the relationship between the amount of input RNA and the number of identified clonotypes from the TRA **(A)** and TRB **(B)** loci across different PCR amplification programs, namely, 18 and 15cycles (18/15), 21 and 18-cycles (21/18) and lastly 24 and 21 cycles (24/21). **(C)** and **(D)** show the relationship between RNA quality, as measured by RIN^e^ score, and the number of TRA **(C)** and TRB **(D)** clonotypes across different PCR amplification programs. **(E), (F)** and **(G)** represent the relationship correlation in the abundance of different TRAV gene segments when comparing libraries prepared using 100 ng to libraries prepared at 200 ng **(E),** libraries prepared using 100 relatives to libraries prepared using 300 ng **(F),** and finally, libraries prepared using 200ng of RNA relative to libraries prepared using 300 ng of RNA **(G).** The same relationship is depicted in **(H), (I),** and **(J)** but for different TRAJ genes, in **(K) (L)**, and **(M)** but for TRBV genes, and lastly in **(N), (O)** and **(P)** for different TRBJ.

After studying the effect of RNA quantity on the sensitivity of TCR-Seq, we examined the effect of RNA quality on the number of identified clonotypes, as well as the interplay between RNA quality and PCR cycle optimization. By increasing the PCR cycles the number of identified clonotypes could be increased, however, the magnitude of increase is a function of RNA quality (**Fig. 2C-2D**). Specifically, samples with low quality (RINe<6) generally yield a smaller number of clonotypes relative to samples with higher RNA quality, nonetheless, increasing PCR cycles in these low-quality samples led to a marginal increase in the number of clonotypes identified. On the other side, increasing PCR cycles in samples with higher RNA quality led to a substantial increase in the number of identified clonotypes, mainly, in samples with low quantities as discussed above. These findings imply that optimizing the amplification strategy is a powerful tool to increase clonotype identification for samples with low quantity and to a lesser extent lower quality.

### The input quantity and the amplification strategy do not bias V and J gene frequency estimation by TCR-Seq

After investigating the impact of RNA quantity and the amplification strategy on the number of identified clonotypes by TCR-Seq, we aimed to study the effect of these variables on estimating the frequency of different V and J genes in the T cell repertoire of different samples. Across the different amplification strategies and RNA quantities, TCR-Seq generated highly consistent frequency estimates of the TRAV genes (**Fig. 2E-2G**), TRAJ genes (**Fig. 2H-2J**), TRBV genes (**Fig. 2K-2M**) and TRBJ genes (**Fig. 2N-2P**). Interestingly, these estimates were more stable for the J genes of the TRA and TRB loci, *i.e*. TRAJ and TRBJ genes, relative to the TRAV and TRBV genes. The stability of frequency estimates for the J-genes can be attributed to the structure of the amplicon where the entire J-gene is sequenced while only a part of the V-gene is sequenced making V-gene calling a harder problem than J-gene calling. Nonetheless, our results indicate that neither the RNA quantity nor the amplification strategy has a meaningful impact on the ability of TCR-Seq to estimate the gene frequency of the different V and J genes in the T cell repertoire.

Subsequently, we wanted to investigate the impact of RNA quality, quantity, as well as PCR amplification on the estimated diversity of the T cell repertoire. For libraries prepared using the 18/15 amplification program, increasing the RNA quantity was associated with a significant increase in the estimated diversity of the repertoire as measured using the Shannon diversity index **(Fig. 3A, 3D**). This effect was evidenced in the repertoires of the TRA (**Fig. 3A**) and the TRB (**Fig. 3D**), especially between libraries prepared from 100 ng relative to libraries prepared from 200 or 300 ng of RNA. Among libraries prepared from 200 and 300 ng of RNA, there was no significant increase in diversity, suggesting that libraries prepared from 100 ng using this PCR amplification program severely underestimated the diversity of the sample, regardless of the RNA quality of these samples. Importantly, increasing the number of PCR program to 21 and 18 or even to 24 and 21 cycles diminishes differences in the diversity of libraries prepared using different RNA quantities for both the TRA (**Fig. 3B-3C**) and TRB (**Fig. 3E-3F**) repertoires. After that we investigated the impact of RNA quality on the estimated diversity of the library, across the two loci and the three PCR amplification programs, RNA quality was positively associated with an increase in Shannon diversity (**Fig. 3G-3L**). These results confirm our previous observation that while increasing PCR cycles leads to an increase in clonal richness for samples with low RNA quantity, it has a minimal or low effect in increasing the richness for samples with low RNA quality.

**Figure 3.**
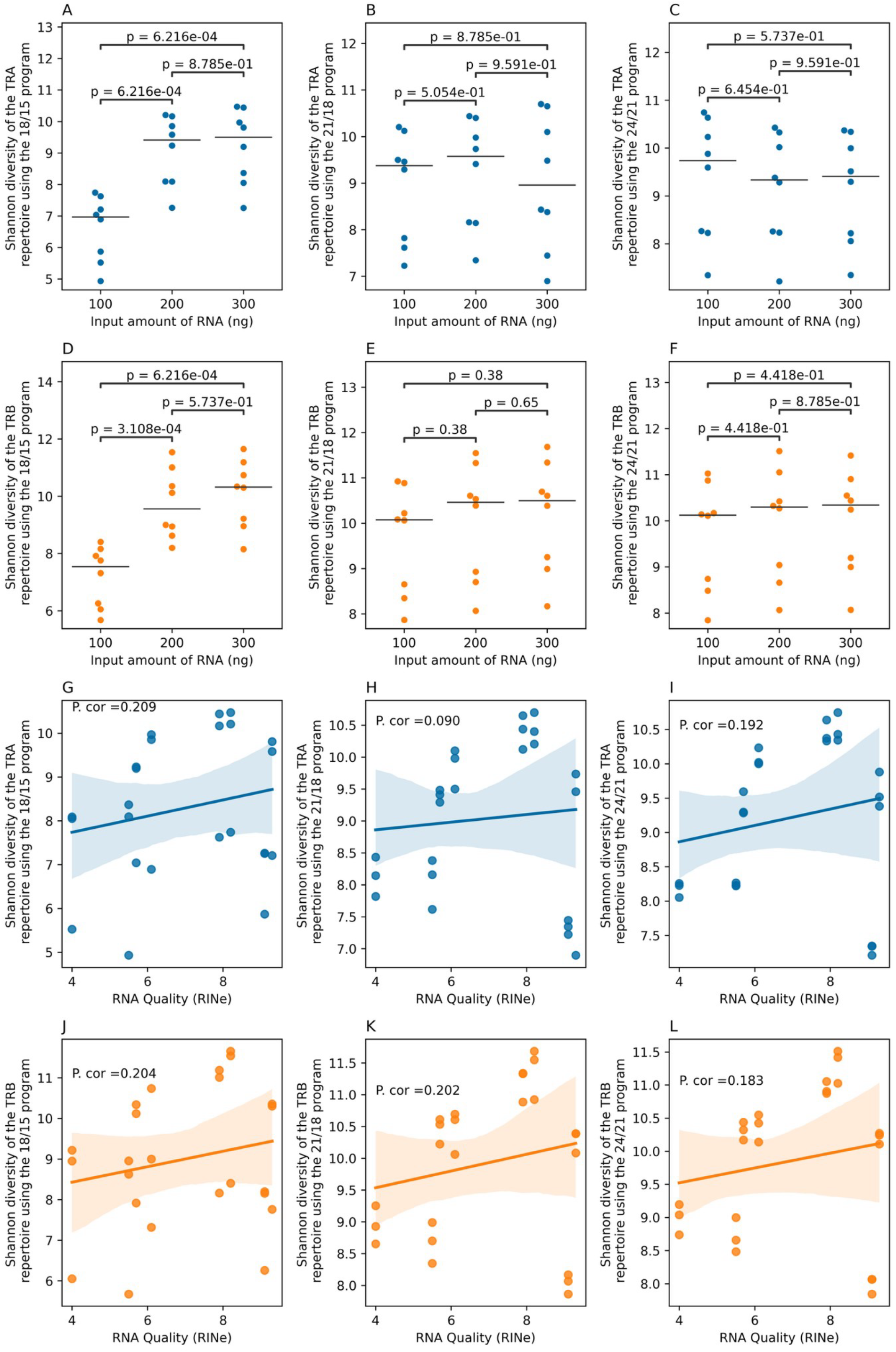
The impact of changing the input quantity and quality as well as the PCR cycles on Shannon diversity. **(A), (B)** and **(C)** show the impact of increasing the PCR cycles program on the estimated Shannon diversity of T cell receptor alpha chain (TRA) across different input quantities, **(A)** shows the diversity after using 18 cycles for PCR1 and 15 cycles for PCR2, **(B)** shows the results using 21 cycles for PCR1 and 15 cycles for PCR2 and lastly **(C)** depicts the diversity of samples processed with 24 cycles in PCR1 and 21 cycles in PCR2. Similarly **(D), (E)**, and **(F)** show the impact of increasing the input quantity on Shannon diversity across different amplification programs with **(D)** showing the results after 18 cycles for PCR1 and 15, **(E)** 21 and 18 cycles and **(F)** 24 and 21 cycles. **(G), (H)** and **(I)** show the relationship between input RNA quality and the estimated Shannon diversity of the TRA chain across different PCR amplification programs with **(G)** showing the results using the 18/15 program, **(H)** the 21/18 program, and **(I)** the 24/21 program. **(J), (K)**, and **(L)** show the correlation between RNA quality and the diversity of the TRB chain across different amplification programs, **(J), (K),** and **(L)** depicts these correlations at the 18/15, 21/18, and 24/21 program, respectively.

### RNA quality and quantity greatly influence the ability of TCR-Seq to identify changes in the T cell repertoire of tissues

Although TCR-Seq is commonly conducted on blood-based biological materials, *e.g.* PAXgene tubes and PBMCs, performing TCR-Seq on biopsies can provide insights into the local T cell repertoire, for example, tissue-resident T cells. To quantify the impact of RNA quantity and quality on shaping the results of TCR-Seq conducted on tissue biopsies, we performed TCRSeq on 26 biopsies collected from different sections of the colon and rectum. These samples had a median quality of ~ 3.75 as measured by the RIN^e^ score and a median concentration of ~100.7 ng/μL (**Fig. S7**). Similar to PAXgene-tubes derived samples shown above, the quality and quantity of RNA played a pivotal role in determining the ability of TCR-Seq to estimate the clonal richness and the diversity of T cell repertoires in tissues. Indeed, for the majority of samples, the T cell repertoire could not be profiled using 300ng of RNA as library preparation and sequencing failed. To mitigate this problem, we performed parallel cDNA synthesis in which multiple cDNA-synthesis reactions were prepared from the same biological material if there was enough material to support this approach. Subsequently, these cDNA libraries were pooled together prior to target amplification in PCR1 which enabled us to increase the amount of input from ~300ng (1X) up to ~1800ng (6X) of cDNA.

This approach increased the estimated clonal richness, *i.e.* the number of identified unique clonotypes, illustrating the impact of the input quantity on shaping TCR-Seq results. Consequently, we were interested in investigating the effect of RNA quality on shaping the identification results at high quantities of input (6X the amount of input RNA). RNA quality strongly correlated with the clonal richness (**Fig. 4A-4B**) and the diversity of the repertoire as defined by the Shannon-diversity index (**Fig. 4C-4D**), illustrating the strong impact of RNA quality that cannot be compensated by increasing the amount of the input RNA, at least within the limits tested here.

**Figure 4.**
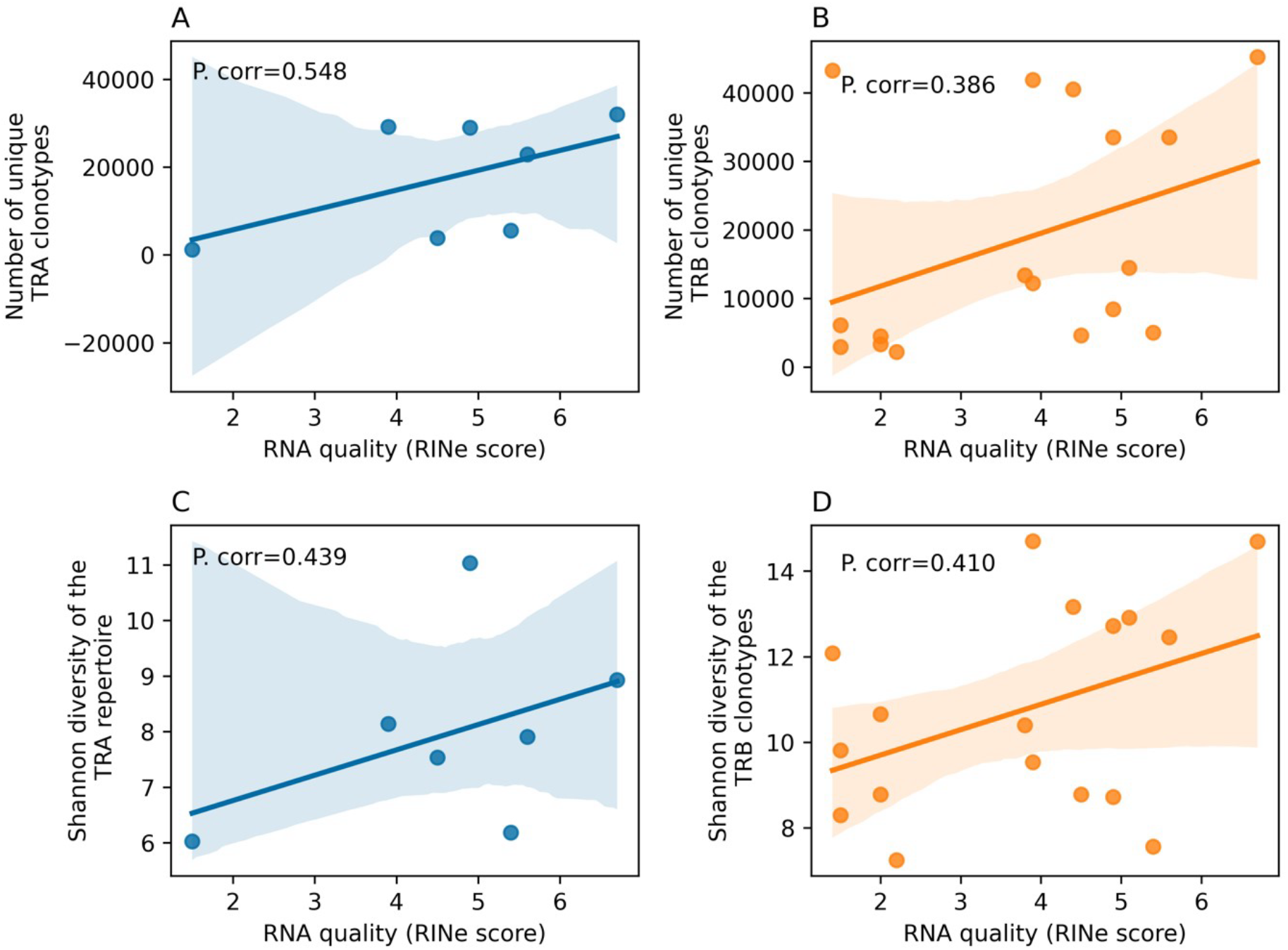
The relationship between RNA quality as defined by the RIN^e^ score and the clonal richness and clonal diversity across the repertoire of TRA and TRB clonotypes. **(A)** and **(B)** depict the strong positive correlation between RNA quality and the number of identified unique TRA **(A)** and TRB **(B)** clonotypes. Similarly, **(C)** and **(D)** illustrate the strong positive correlation between the quality of input RNA and the estimated repertoire diversity using the Shannon diversity index for TRA **(C)** and TRB **(D)**.

### The nucleic acid extraction protocol has a minor impact on repertoire profiling results

After studying the interplay between RNA quality, RNA quantity, and the amplification strategy, and their influence on TCR-Seq results, we were interested in investigating the impact of RNA extraction method. Thus, we collected 4 PAXgene tubes from a single healthy donor at a single timepoint, then extracted RNA using two robotic extraction methods, namely, QIAGEN QIAcube connect with in vitro diagnostics kits (IVD1) (n=2 tubes) and QIAGEN QIAcube connect MDx (MDx2-IVD2) (n=2 tubes), which both generated high quality RNA (RINe>9). Subsequently, 300ng of RNA were used for profiling the TRB repertoire of these four biological replicates using the same PCR amplification program. We primarily focused here on the TRB repertoire because of its higher diversity relative to the TRA repertoire, hence, it will be more sensitive to any differences introduced by the different RNA extraction methods. After pooling, deep sequencing was conducted on the generated libraries, generating >20 million reads per library. Lastly, the generated sequencing results were analyzed using MiXCR as discussed previously (**Material and Methods**). These four repertoires showed minimal overlap with each other (**Fig. 5A**), more importantly, we did not observe any significant clustering or increased similarity among replicates sharing the same RNA-isolation method (**Fig. 5B**). This suggest that the variability introduced using the RNA extraction protocol is either smaller than the biological variability present among the different replicates and/or smaller than technical variability introduced by different steps in the protocol. Despite the limited overlap in the identified clonotypes, the estimated frequencies of the different TRBV genes in each of the samples, strongly correlated among the four samples (**Fig. 5C-5H**). Indicating that neither of the methods results in a systematic enrichment or depletion of clonotypes belonging to a particular TRBV genes. Thus, within the two tested systems, no detectable impact for the extraction protocol on the identified repertoire was observed.

**Figure 5:**
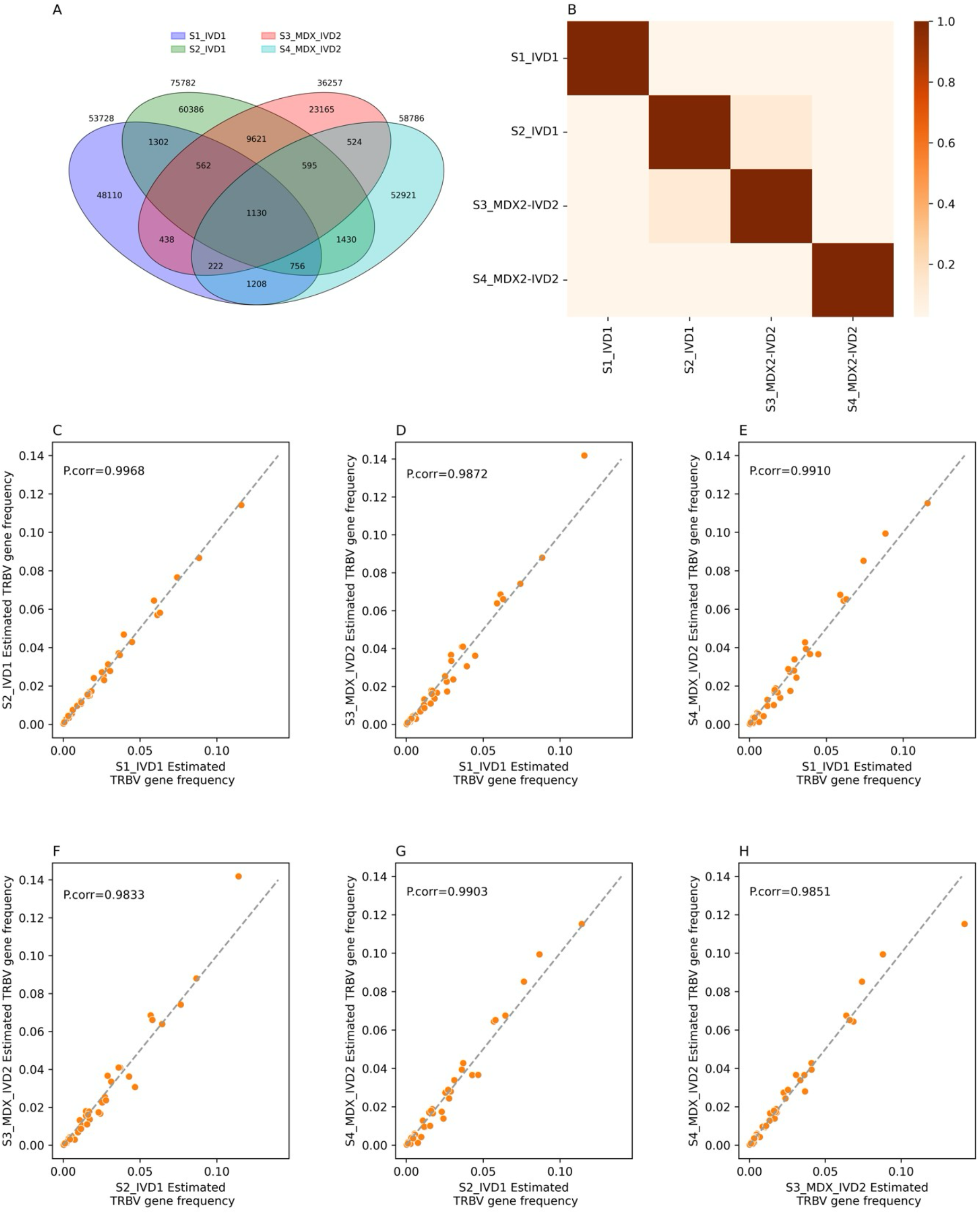
RNA extraction protocol has a minimal impact on the identified TCR repertoire. (**A**) depicts the overlap among the four replicates in the term of clonotypes which are defined as unique combination between V-genes, Jgenes and CDR3 amino acid sequences. Across all panels, S1_IVD1 and S2_IVD1 refer to the first and second replicates extracted using the IVD1 system while S3_MDX_IVD2 and S4_MDX_IVD2 donate replicate 3 and replicate 4 that have been extracted using the MDX IVD2 robotics system. (**B**) illustrate the overlap among the samples using Jaccard-index, without a clear clustering based on the RNA extraction framework. (**C**)-(**H**) show the strong correlation between estimated TRBV gene frequency among different pair of the four replicates, *e.g.* (**C**) depicts the correlation in the TRBV gene frequency between S1_IVD1 and S2_IVD2, while (**D**) depicts the correlation between S1_IVD1 and S3_MDX_IVD2.

### DNA quantity plays a pivotal role in shaping clonotype identification from gut biopsies in a loci-specific manner

After analyzing the results of RNA-based assays, we aimed to examine the results of DNAbased assays. To this end, we profiled the full immune repertoire, that is, the T and B cell repertoire, of six different tissue biopsies using three different amounts of DNA, specifically, 40, 80, and 120 ng (**Material and methods**). Increasing the amount of DNA was significantly associated with an increase in clonal richness across some loci, such as TRA, TRB, and TRG while it had a minor-to-no effect on the clonal richness of other loci, such as TRD, IGH, and IGK (**Fig. S8**). Suggesting, that the diversity of these loci in the sample was on average low enough that ~40ng of DNA was enough to capture most of the unique clonotypes derived from these loci. To follow this further, we calculated the Shannon-diversity index (**Material and methods**) for all loci across the different input amounts. The diversity calculations recapitulated the clonal-richness results and correlated positively with increasing the amount of input DNA across all loci except TRD where increasing the amount of input DNA did not affect diversity (**Fig. S9**). Lastly, we looked at Simpson’s evenness (**Material and methods**) which yielded a pattern resembling the results of Shannon diversity, where increasing the amount of input DNA results in a decrease in the evenness across all loci except TRD loci (**Fig. S10**). Thus, similar to RNA increasing the amount of DNA used for library preparation has a major influence on shaping the results of TCR-Seq experiments such as clonal richness, estimated diversity, and evenness.

## Discussion

In this study, we investigated the impact of different technical variables on TCR-Seq results. A key finding throughout our analyses is the crucial role of RNA quality, which strongly correlated with the richness and diversity of the identified repertoire. Although increasing the quantity of RNA and number of PCR cycles tend to enhance the detection of clonotypes within a sample, these gains are bounded by the RNA quality. Hence, to ensure deep and comprehensive profiling of the T cell repertoire, maximizing RNA quality is essential whenever possible. Although RNA quantity also strongly correlated with repertoire richness and diversity, it was relative to quality, easier to optimize by adjusting the amplification program.

Profiling the repertoire of tissue biopsies was more challenging than profiling peripheral blood, mainly due to the differences in cellular composition. Blood contains a relatively high number of T cells compared to most other tissues, such as colon biopsies. This implies that when using the same amount of RNA or DNA, blood samples typically yield more T cell-derived material. As a consequence, larger amounts of RNA are needed to profile the T cell repertoire from tissue biopsies at the same resolution as blood. A potential solution to this problem is to perform multiple cDNA reactions in parallel prior to pooling them for the initial PCR reaction. Although this might be a practical solution, our results suggest the need to develop amplification kits and protocols that are tailored to process and analyze tissue biopsies. For example, increasing the amount of DNA from 40 to 120 ng significantly increased the clonal richness of the TRA, TRB and TRG repertoires, but not the TRD repertoire, which has limited diversity that was captured using only 40ng of DNA. This highlights the need to adjust the protocols to account for fewer T cells and varying diversity between the different T cell subpopulations present in a tissue, such as the less diverse TRD repertoire and more diverse TRG repertoire.

## Funding

The project was funded by the EU Horizon Europe Program grant *miGut-Health: Personalized Blueprint of Intestinal Health* (ID: 101095470). Additionally, the project received funding from the German Research Foundation (DFG) Research Unit 5042: miTarget (The Microbiome as a Therapeutic Target in Inflammatory Bowel Diseases) along with funding from the DFG Cluster of Excellence 2167 ‘Precision Medicine in Chronic Inflammation (PMI)’. A.K.H.M. is funded by the DFG collaborative research center CRC 1526 ‘Pathomechanisms of Antibody-mediated Autoimmunity (PANTAU) – Insights from Pemphigoid Diseases’. This work was supported by the DFG Research Infrastructure NGS_CC (project 407495230) as part of the Next Generation Sequencing Competence Network (project 423957469). NGS analyses were carried out at the Competence Centre for Genomic Analysis (Kiel).

## Availability of Data

Data are not deposited in a public repository due to data privacy regulations in Norway and our institution. However, data are available upon reasonable request, if the aims of the planned analyses are covered by the written informed consent signed by the participants, pending an amendment to the ethical approvals and a material & data transfer agreement between the institutions.

### Material and Methods Library preparation from RNA and DNA materials

All library preparation methods were conducted according to the manufacturer’s instructions. As discussed in the results section two variables varied among the experiments. First, the starting material which ranged from 100 to 300 ng of RNA for blood-based RNA samples, 300 to 1800 ng of RNA from gut biopsies, and 40 and 120 ng of DNA from biopsies samples.

Second, the PCR amplification program as described in the Material and Methods sections.

### Sequencing approach

Following library preparation, illumine indices were added to each prepared library prior to the multiplexing of these libraries together. After cleaning with magnetic beads, the pooled libraries were diluted to ~2nM and were subsequently submitted to sequencing at the <u>C</u>ompetence <u>C</u>entre for Genomic <u>A</u>nalysis (CCGA). To improve cluster resolution on the flow cell 10-15 PhiX were used with each pool. Sequencing of RNA samples was conducted using NovaSeq 6000 on an S4 flow cells using 150bp paired read sequencing. The number of reads per pool was selected to allow for an average of 5 million reads for TRA and ~5 million reads of TRB libraries. For DNA libraries, sequencing was also done using the same framework, *i.e.* 150bp paired sequencing on NovaSeq 6000 S4 cells. However, because the utilized library preparation kit amplifies the seven immune receptor chains in a single library, each library was aimed for ~10-15 million reads per library.

### Clonotype identification and computational analysis

After sequencing and samples demultiplexing, MiXCR^7^ (v4.6.0 with RepSeq.IO v2.4.0; MiLib v3.4.0 and the Built-in V/D/J/C library utilizing repseqio. v4.0) was used for clonotype identification for each sample. For libraries prepared using the Human TCR RNA Multiplex kit with UMI (MiLaboratories), the analysis was conducted using the ‘milab-human-tcr-rnamultiplex-cdr3’ preset. On the other side, for libraries prepared using the 7genes DNA multiplex kits (MiLaboratories), the analysis was conducted using the ‘milab-human-dna-xcr7genes-multiplex’ preset. After exporting clonotypes from each library, non-productive clonotypes, i.e. clonotypes that do not encode a functional immune receptor chain, were removed. Subsequently, custom code was used to merge V(D)J rearrangements that utilize the same V and J genes and produce the same CDR3 sequence at the protein level together. During the merging step, the count and fractions of UMIs and reads were respectively added together, to produce a final UMI and read count as well as a reaction for each unique combination of V, J genes, and CDR3 sequence in each library. Subsequently, samples were combined into a single table and custom Python code was used for running all the analyses shown above.

## Notes

### Competing Interest Statement

The authors have declared no competing interest.

