## Supplementary figures for "Quantifying The Impact of Bulk TCR-Seq Methodological Choices on The Profiled T Cell Repertoire"

### Supplementary figure

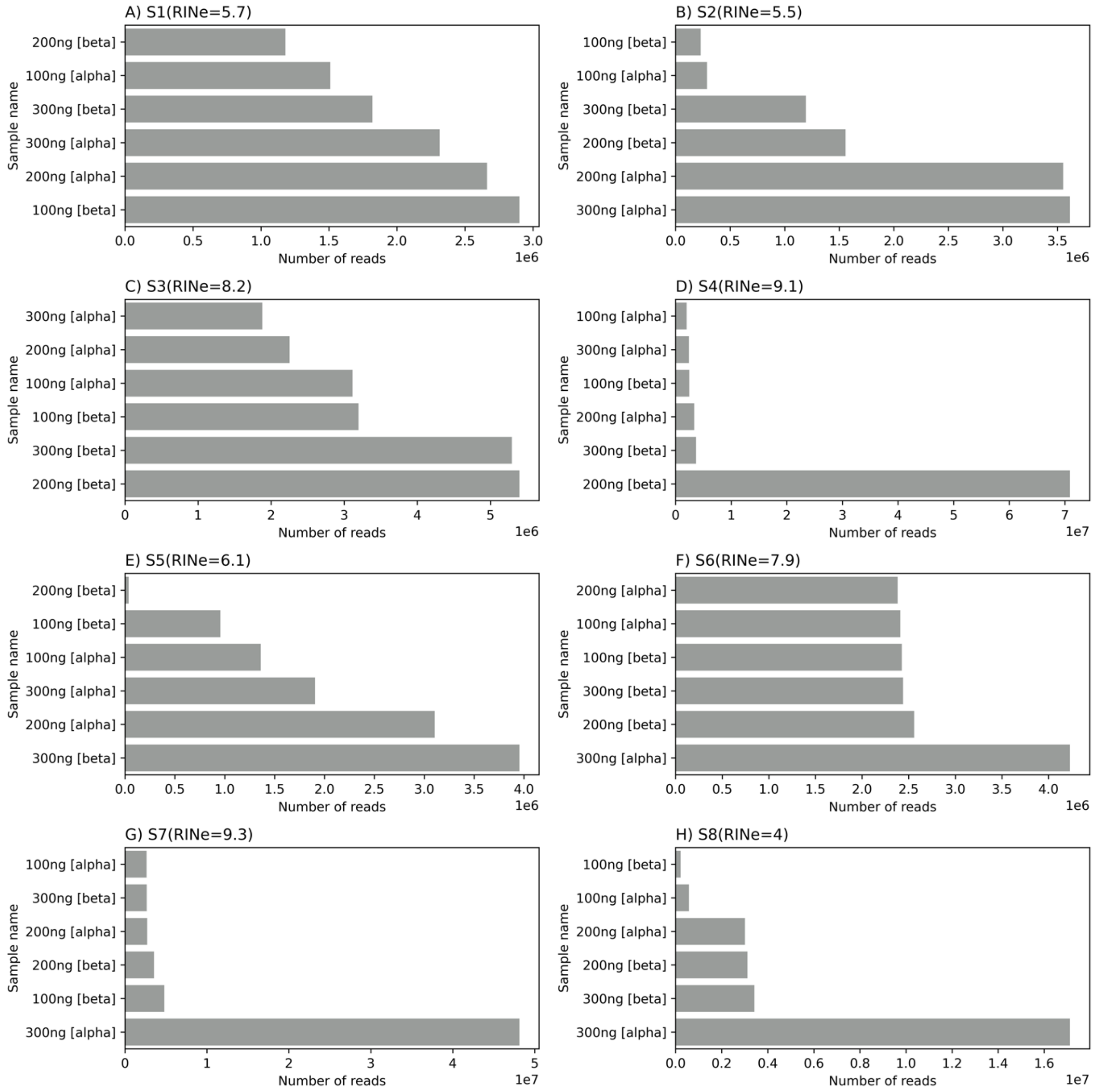


**Figure S1:** *The number of reads derived from the eight different samples utilized in the current study.* For each panel **(A-H)**, the number of reads derived from the different libraries prepared using different amounts of RNA is shown.

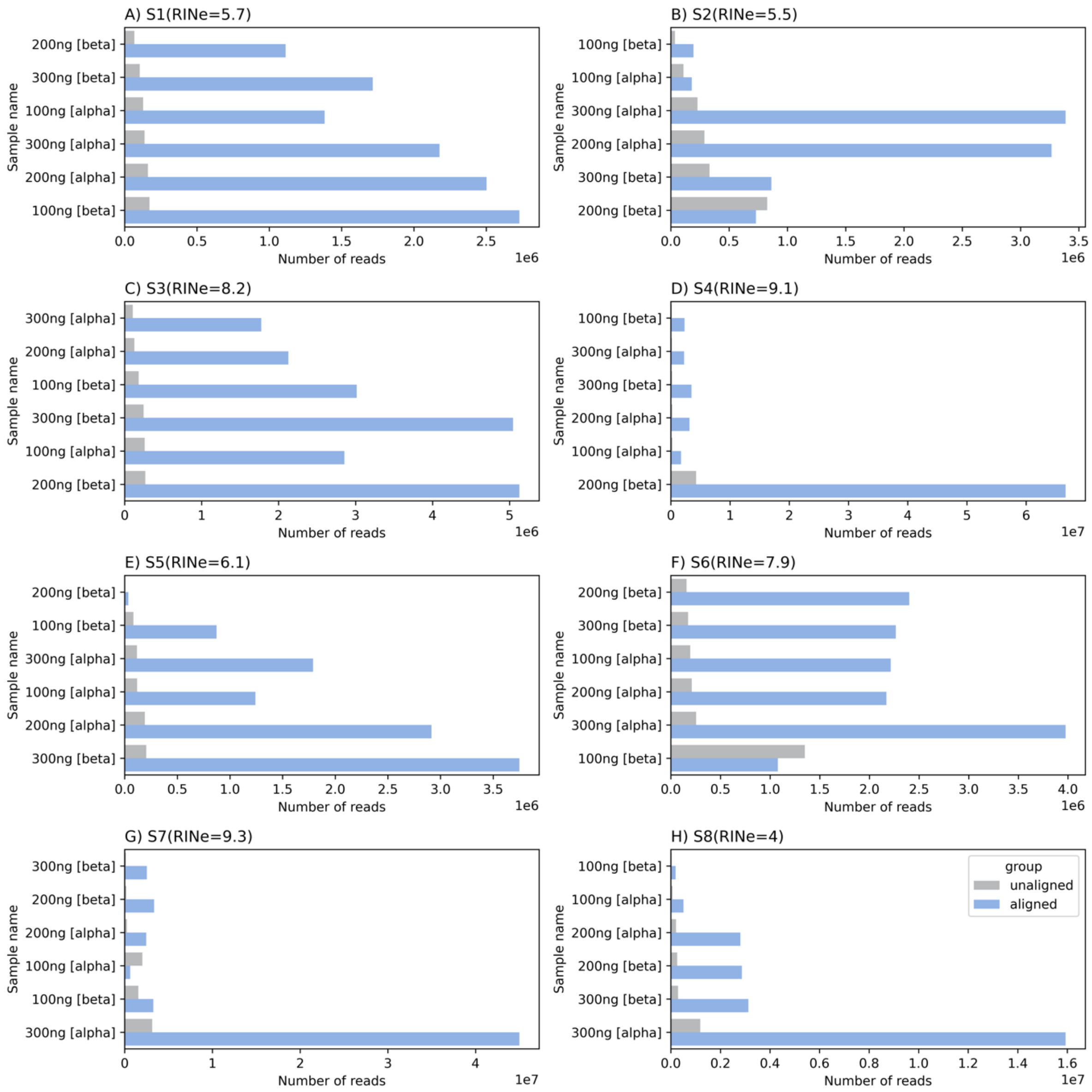


**Figure S2:** *The number of aligned and unaligned reads from libraries prepared in the current study. Panels (**A-H**) represent the eight different samples used in the current study.* For all panels, the y-axis represents the different libraries prepared from each sample and the x-axis represents the number of reads.

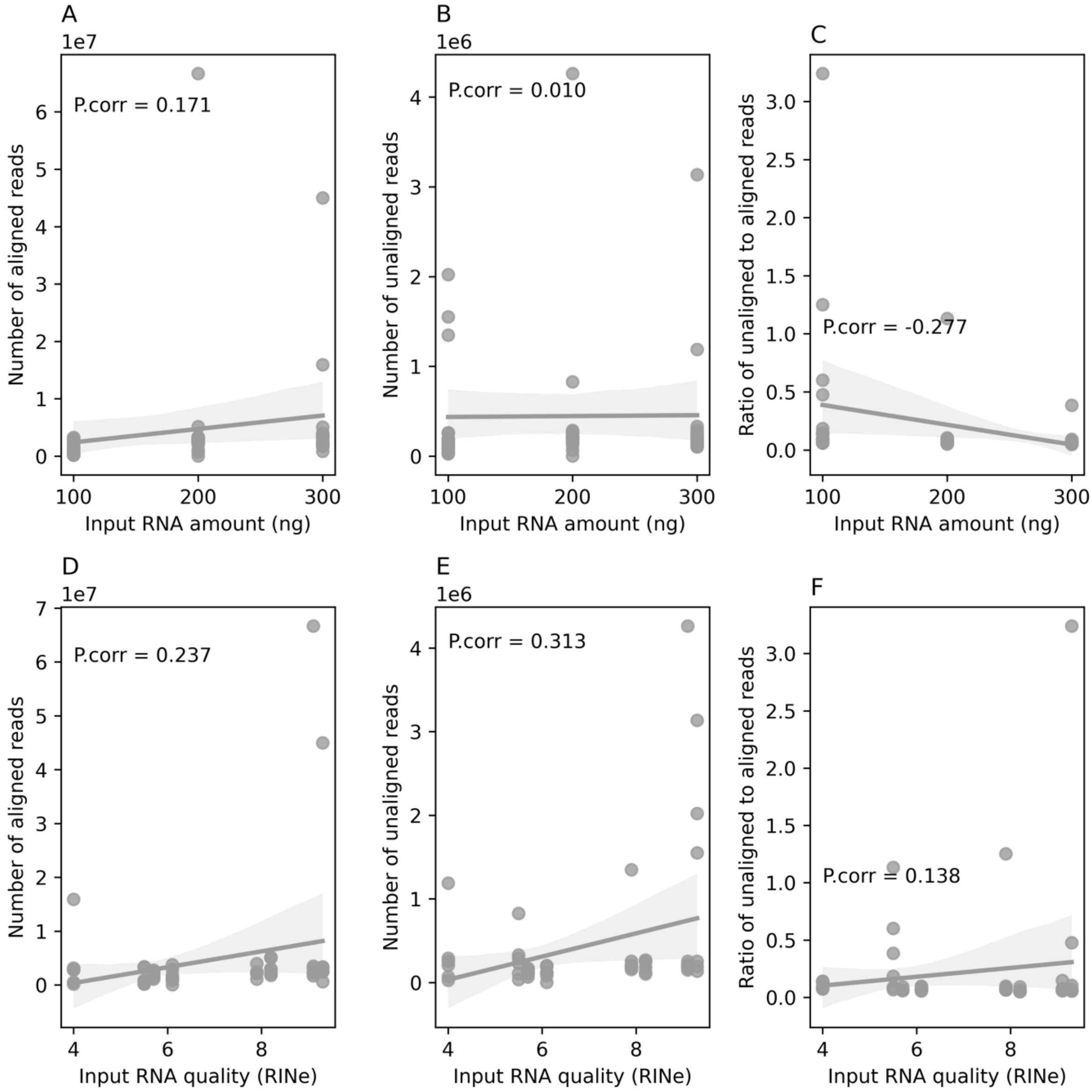


**Figure S3:** *The relationship between RNA quantity and quality and the number of aligned and unaligned reads per library*. **(A)** depicts the relation between increasing the amount of input RNA and the number of aligned reads. **(B)** shows the relationship between increasing the RNA quantity and the number of unaligned reads. **(C)** illustrate the negative correlation between increasing the amount of RNA and the ratio of unaligned to aligned reads per library. **(D)** shows the relationship between RNA quality and the number of aligned reads. Similarly, **(E)** shows the relationship between RNA quality and the number of unaligned reads. Lastly, **(F)** shows the relationship between RNA quality and the ratio of unaligned to aligned reads.

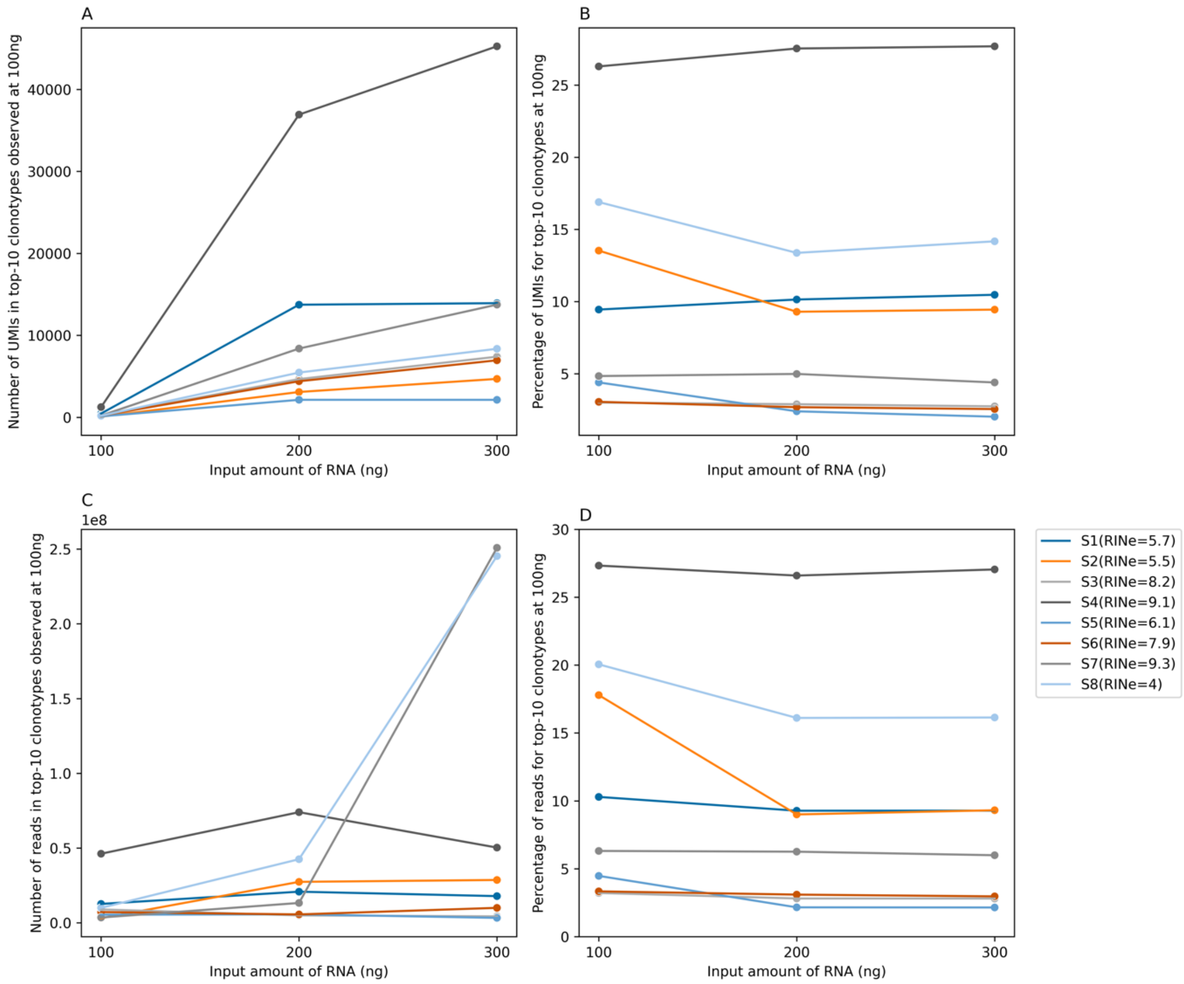


**Figure S4:** *The relationship between the starting amount of RNA and the bias in detecting and quantifying highly expanded clonotypes.* **(A)** depicts the relationship between increasing the starting RNA amount on the number of UMIs that top-10 clonotypes detect at 100 ng by increasing the amount to 200 and 300 ng. **(B)** shows the same relationship but after normalizing with the total number of UMIs and plotting the percentage of UMIs derived from this set of top-10 clonotypes as a percentage instead of absolute counts as shown in **(A)**. **(C)** and **(D)** illustrate the relationship between increasing the amount of RNA and the number of reads top-10 clonotypes have as absolute count (C) and as a percentage of total reads **(D)**.

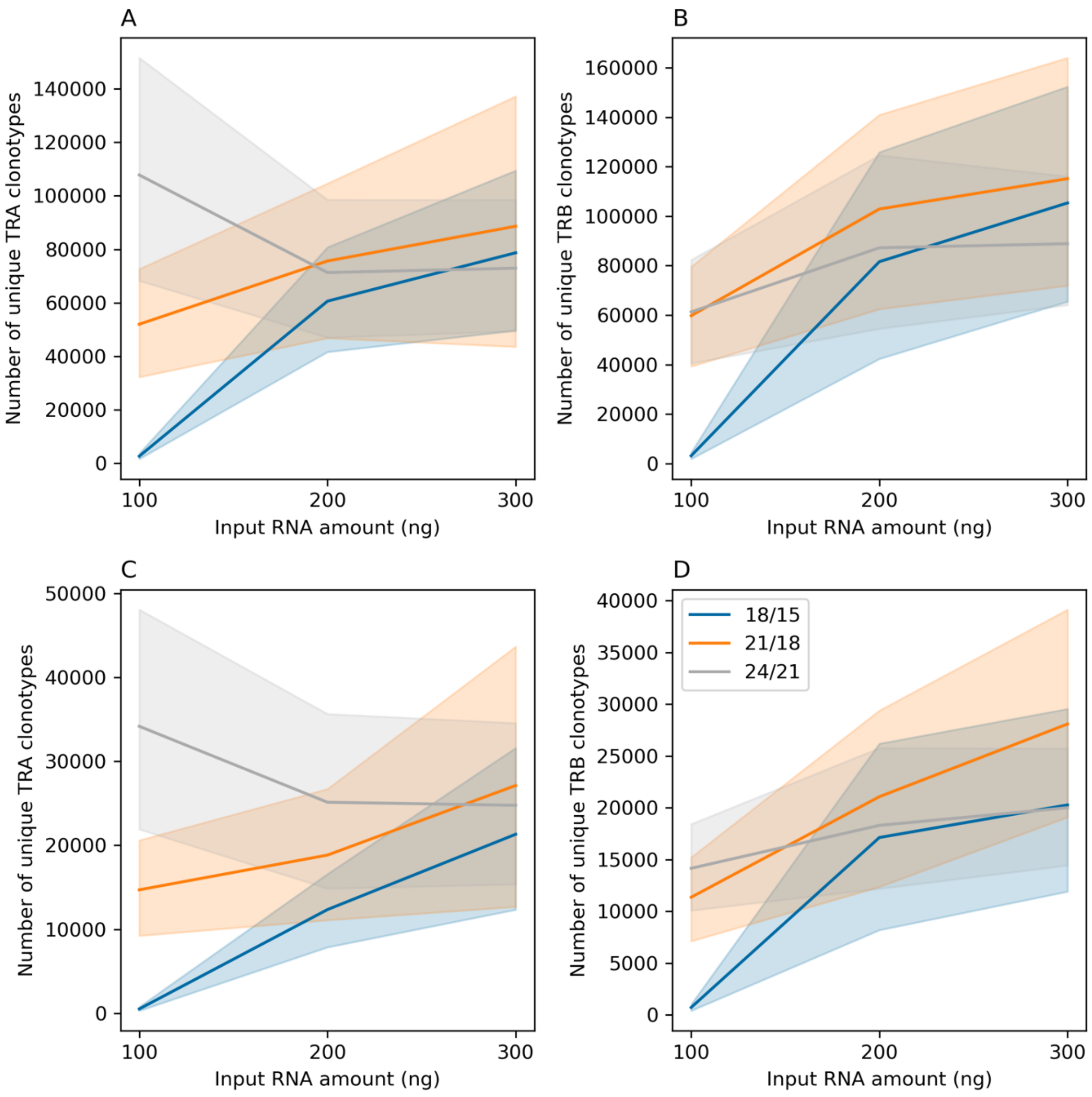


***Figure S5**: The impact of increasing the amount of RNA and the number of identified clonotypes from the T cell receptor alpha chain (TRA) and T cell receptor beta chain (TRB) using three different PCR amplification strategies, namely, 18 cycles in the first PCR reaction followed by 15 cycles in the second PCR reaction (18/15) or 21 cycles followed by 18 cycles (21/18) and lastly, 24 cycles followed by 18 cycles (24/18).* **(A)** and **(B)** duplicates the relationship between the amount of RNA used for preparing the library and the number of clonotypes identified under the three amplification strategies after removing clonotypes inferred from only one read. The same relationship is shown in **(C)** and **(D)** but after removing clonotypes supported by less than two UMIs.

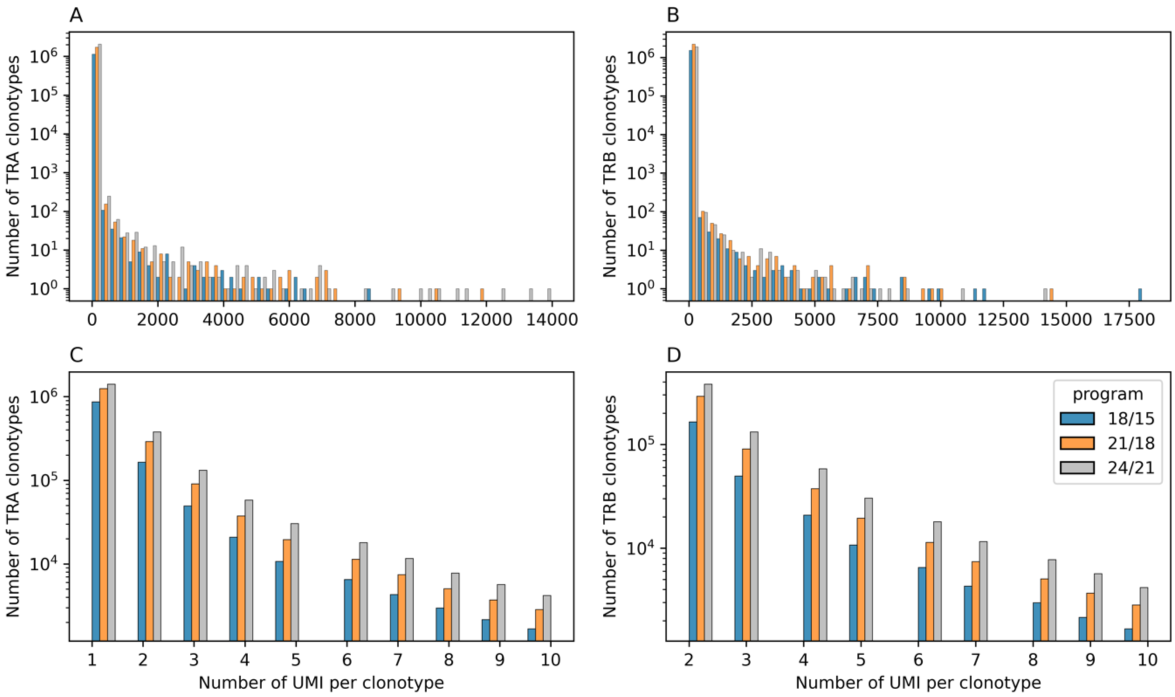


***Figure S6:*** *The number of UMIs per clonotypes*. **(A)** depicts the number of UMIs per each TRA-clonotype identified while **(C)** provides a focused view on TRA-clonotypes having 1-10 UMIs per clonotype. A similar representation for TRB-clonotypes is shown in **(B)** and **(D)**, respectively.

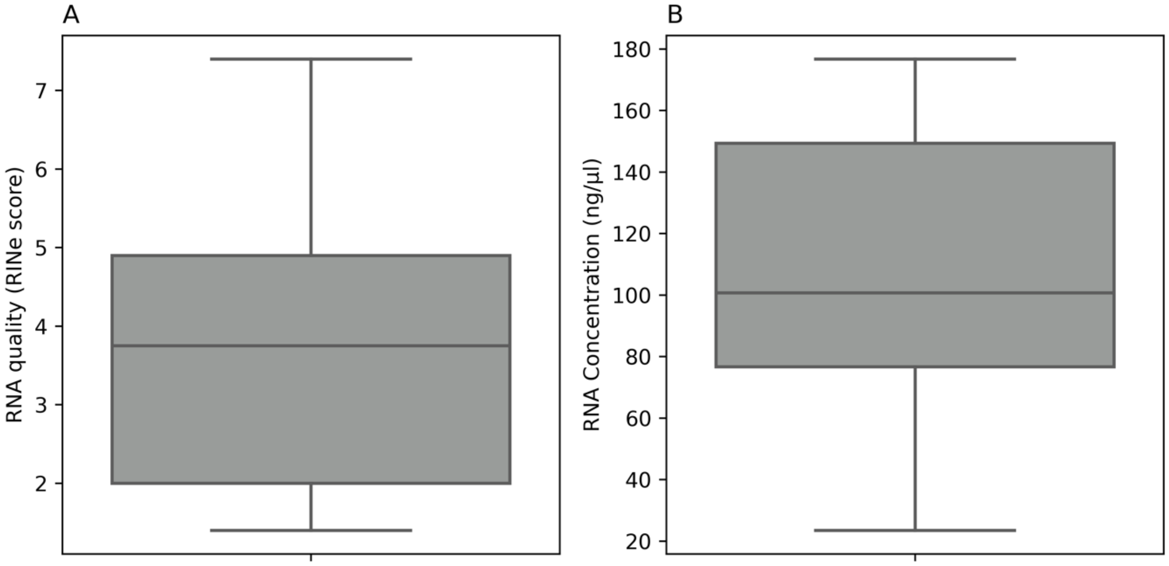

***Figure S7:*** *The quality and concentration of RNA samples isolated from tissue biopsies.* **(A)** shows the distribution of RNA quality as measured by the RINe score while **(B)** shows the RNA concentration.

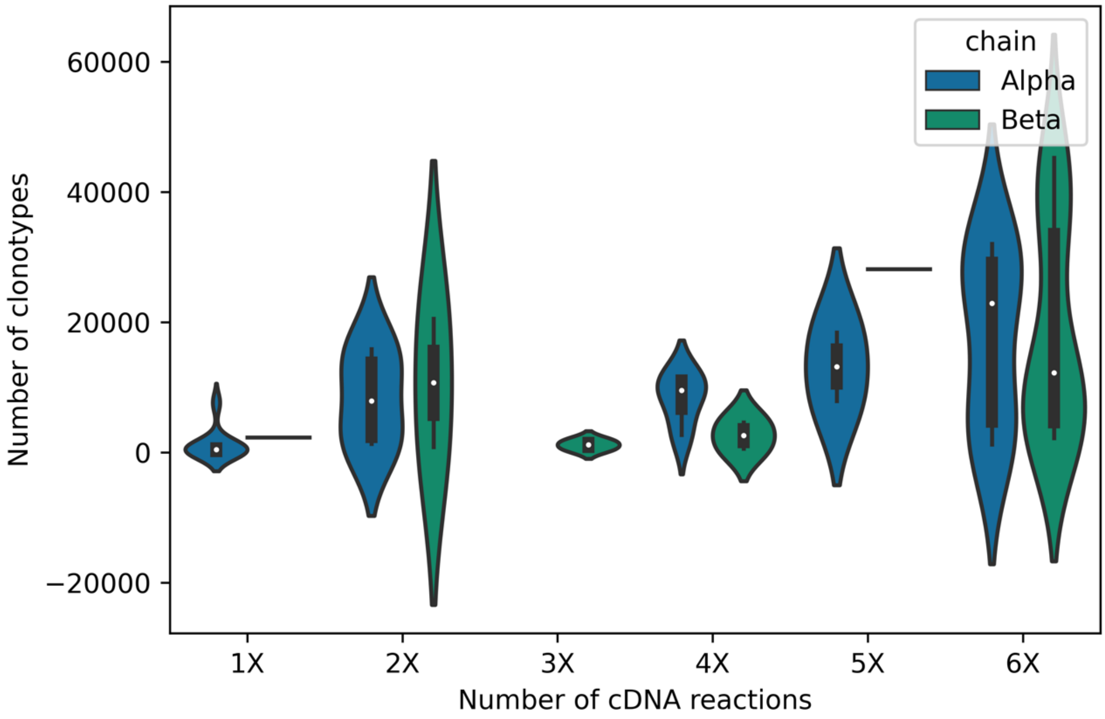


**Figure S8:** *The correlation between increasing the number of cDNA reactions pooled together prior to target amplification in PCR1 and the number of unique clonotypes identified*.

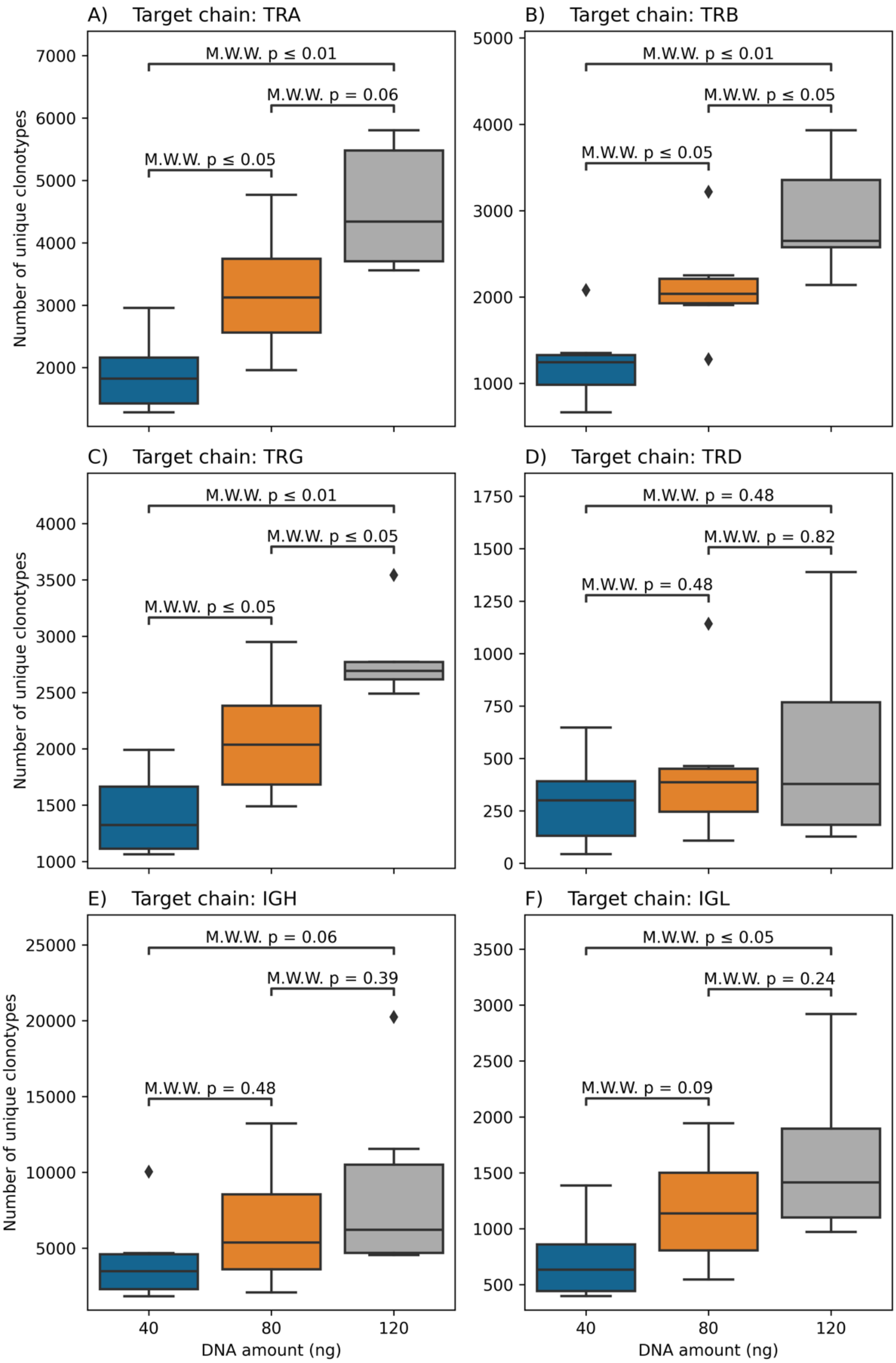


**Figure S9:** *The relationship between increasing the amount of DNA used for library preparation and the number of clonotypes identified from different immune receptor genes*. In **(A)** and until **(F), the** Mann-Whitney-Wilcoxon test was used to compare the number of unique clonotypes, *i.e.* clonal richness, across libraries prepared with different amounts of DNA.

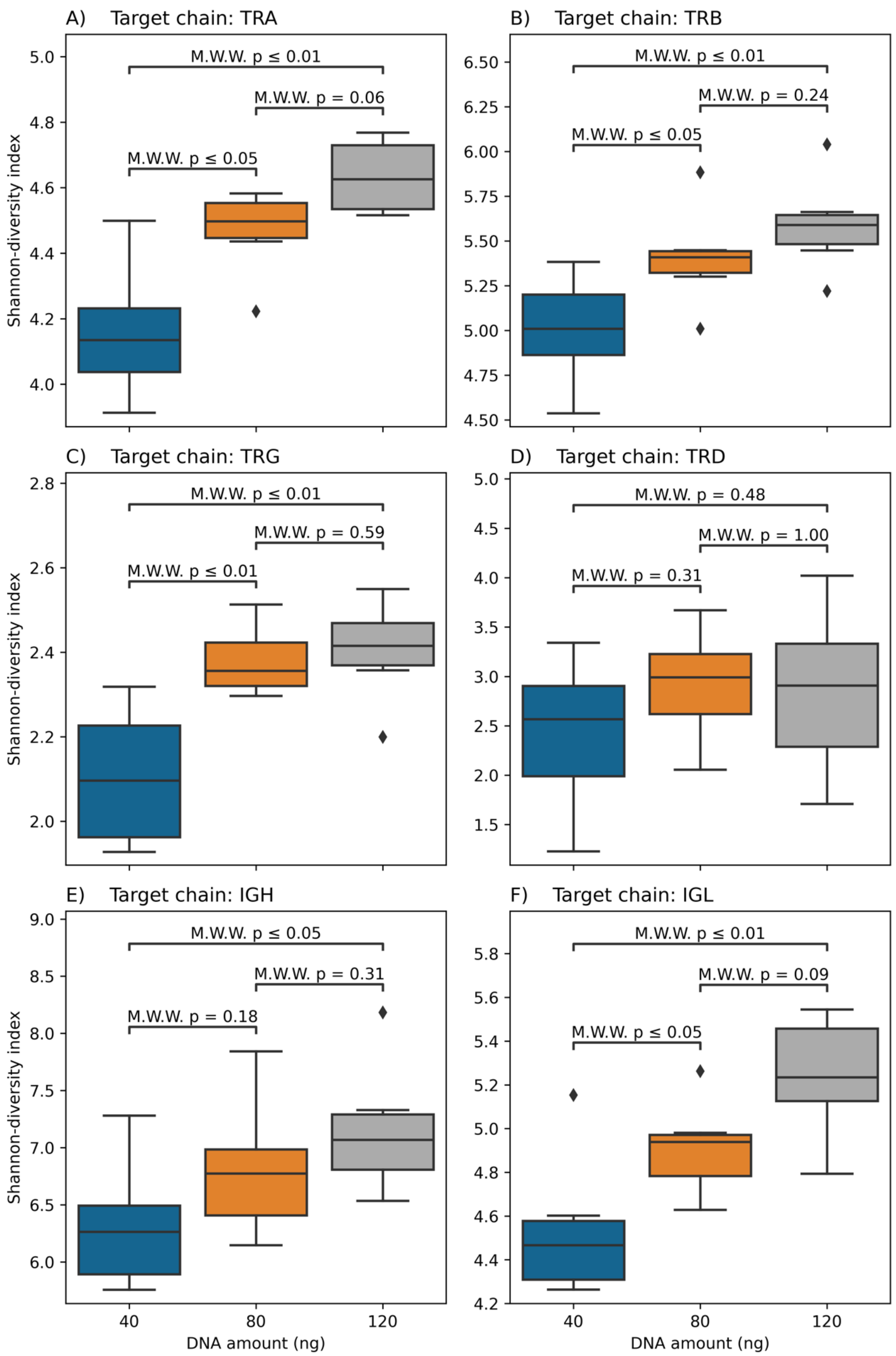


***Figure S10:*** *The relationship between increasing the amount of DNA used for library preparation and the Shannon-diversity index across different immune receptor genes*. In **(A)** and until **(F), the** Mann-Whitney-Wilcoxon test was used to compare Shannon diversity across libraries prepared with different amounts of DNA.

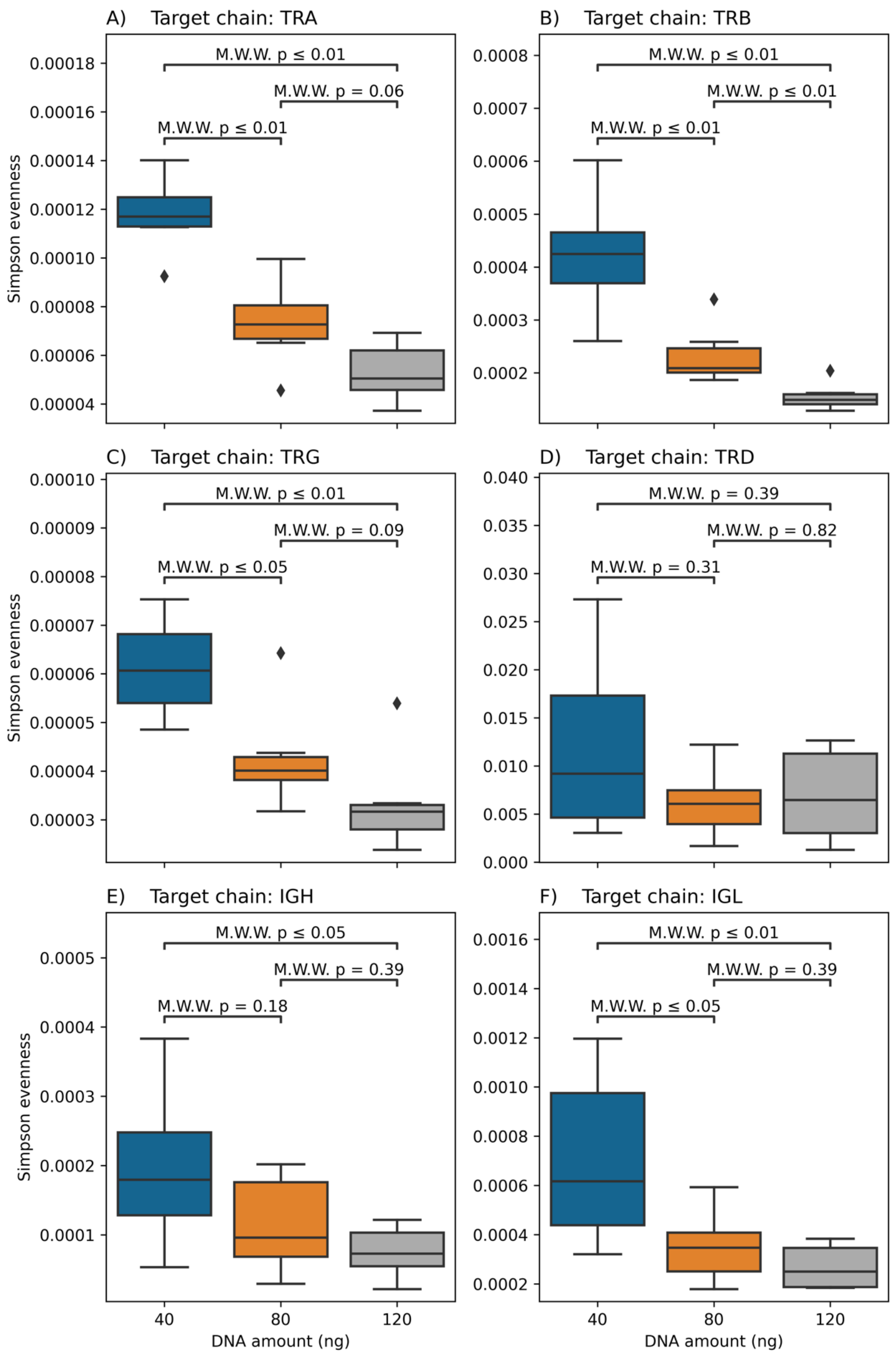


***Figure S11:*** *The relationship between increasing the amount of DNA used for library preparation and Simpson evenness across different immune receptor genes*. In **(A)** and until **(F), the** Mann-Whitney-Wilcoxon test was used to compare Simpson evenness across libraries prepared with different amounts of DNA.
